# Extracellular membrane tubules involved in suberin deposition in plant cell walls

**DOI:** 10.1101/2021.02.02.429332

**Authors:** Damien De Bellis, Lothar Kalmbach, Peter Marhavy, Jean Daraspe, Niko Geldner, Marie Barberon

## Abstract

Suberin is a fundamental plant biopolymer, found in protective tissues, such as seed coats, exodermis and endodermis of roots, the outer layers of stems and roots with secondary growth, as well as in wound-induced tissues. Its presence allows organs to resist various environmental stresses, such as pathogen attack, drought or excessive salt concentrations. Suberin is a mostly aliphatic polyester of long-chain fatty acids and alcohols, often co-occurring with lignin-like polymers in the same cells. Most suberizing cells appear to deposit suberin in the form of lamellae just outside of the plasma membrane, below the primary cell wall. The monomeric precursors of suberin are thought to be glycerated fatty acids, synthesized at the endoplasmic reticulum. However, it has remained obscure how these monomers are transported outside of the cell, where they will be polymerized to form suberin lamellae. Here, we demonstrate that extracellular vesicular-tubular structures accumulate specifically in suberizing cells. By employing various, independent mutational and hormonal challenges, known to affect suberization in distinct ways, we demonstrate that their presence correlates perfectly with root suberization. Surprisingly, no endosomal compartment marker showed any conspicuous changes upon induction of suberization, suggesting that this compartment might not derive from endosomal multi-vesicular bodies, but possibly form directly from endoplasmic reticulum subdomains. Consistent with this, we could block formation of both, suberin deposition and vesicle accumulation by a pharmacogenetic manipulation affecting early steps in the secretory pathway. Whereas many previous reports have described extracellular vesicle occurrence in the context of biotic interactions, our results suggest a developmental role for extracellular vesicles in suberin formation.

**One Sentence Summary:** Suberin lamellae formation is associated with extracellular membrane tubules.

## Main Text

Extracellular vesicles or tubules (EVs) are nanosized membrane-encapsulated structures involved in the secretion of various molecular cargos including proteins, nucleic acids, metabolites and lipids ^1^. A range of vesicles of different size and cellular origin including microvesicles (50-1000 nm) budding from the plasma membrane (PM) and exosomes (50-150 nm) derived from multivesicular body (MVB)-PM fusion are subsumed by the term EV. In mammals, EVs release biomolecules into the extracellular space for targeted intercellular communication. In plants, EVs have been reported early on under various designations (paramural bodies, plasmalemmasomes or boundary structures) and speculated to be associated with cell wall synthesis ^2-4^. In the last decades, EVs have been reported to contain RNAs, defense compounds and signaling lipids and are considered to play a central role in inter-organism communications during defense and symbiosis^5-11^. More recent data implicating EVs in cell wall formation and modification were mostly reported in the context of induced defense responses ^11-14^. It is evident that any type of plant cell wall formation relies on a multitude of secreted molecular building blocks and enzymes for its construction ^15^, yet little has been reported concerning the role for EVs in general cell wall formation during development. EV containing bodies (referred to as “paramural bodies”) were reported to be increased in vesicle trafficking mutants ^16,17^, but it remains largely unknown whether EVs are involved in the regular deposition of cell wall polymers during plant growth and development.

Suberin is a major secondary cell wall formation in plants. In young, primary roots, it occurs exclusively in the endodermis ^18^. By using various genetic, as well as hormonal perturbations of suberin deposition, we demonstrate a strict association of bodies containing extracellular vesicular-tubular structures (EVBs) with suberin deposition in the cell wall. Moreover, we demonstrate that inhibition of the early secretory pathway interferes with both EVB formation and suberin accumulation, suggesting that EVBs are required for the transport of suberin precursors or biosynthetic enzymes to the apoplast and formation of this major secondary cell wall in plants.

In our efforts to understand endodermal differentiation, we performed several genetic screens for endodermal barrier mutants ^19,20^. One screen identified the *lord of the rings 2* mutant (*lotr2/exo70a1*), displaying a fully delocalized Casparian strip membrane domain and an absence of a Casparian strip (CS)^20^. When analyzed at the ultrastructural level, we found a high accumulation of large, vesicle-containing membrane bodies fused with the PM, exclusively in endodermal cells (Fig. 1A, Supplementary Fig. 1A). Such bodies were not observed in wild-type at the same stage of differentiation. We initially thought of this phenotype as a direct consequence of a defective exocyst action in the *lotr2/exo70a1* mutant. However, when we investigated other, unrelated CS-defective mutants, such as *esb1 (enhanced suberin 1)* or *casp1_casp3 (casparian strip membrane domain protein 1* and 3), we found that they equally displayed many such large PM-contiguous bodies, specifically in endodermal cells (Fig. 1B, Supplementary Fig. 1A,B), indicating that the enhanced presence of these bodies is caused by a defective CS and is not a direct consequence of a defective exocyst in the mutant. A 3D reconstruction using FIB-SEM in *lotr2/exo70a1* illustrates the high number and broad distribution of these bodies in an endodermal cell (Fig. 1C, Supplementary Fig. 1C,D, Supplementary Movie 1). These large bodies containing extracellular, vesicular-tubular structures (EVBs) are between 300 and 900 nm in size (Supplementary Fig. 1E) and their extracellular vesicles or tubules were found to be of varying density (Fig. 1A,B, Supplementary Fig. 1B,F) and between 10 and 100 nm in diameter on sections (Supplementary Fig. 1G). We then undertook tilt-series tomographic reconstructions of single bodies in *lotr2/exo70a1*, as well as a wild-type root (where these bodies occur later during endodermal development, see below), in order to understand whether the vesicle structures observed on sections are indeed vesicles, or rather transversally sectioned tubules. The tomograms revealed that the extracellular membrane structures are actually part of a highly branched network of tubules (Fig. 1D; Supplementary Movie 2), with little occurrence of isolated vesicles. Close inspection of a number of tomograms allowed us to identify rare cases in which the outer membrane appeared continuous with an inner tubule, suggesting that the EVBs might be generated by, and are sites of, active evagination (Fig. 1E; Supplementary Movie 3). We also repeatedly observed a very thin electron dense layer at the surface of suberin lamellae in the tomograms (Fig. 1F; Supplementary Movie 4). This fits with a scenario in which the nearby tubules/vesicles fuse with the surface of the suberin lamella, the thin black layer representing the rest of the membrane after fusion with the lipidic lamella surface.

**Figure 1.**
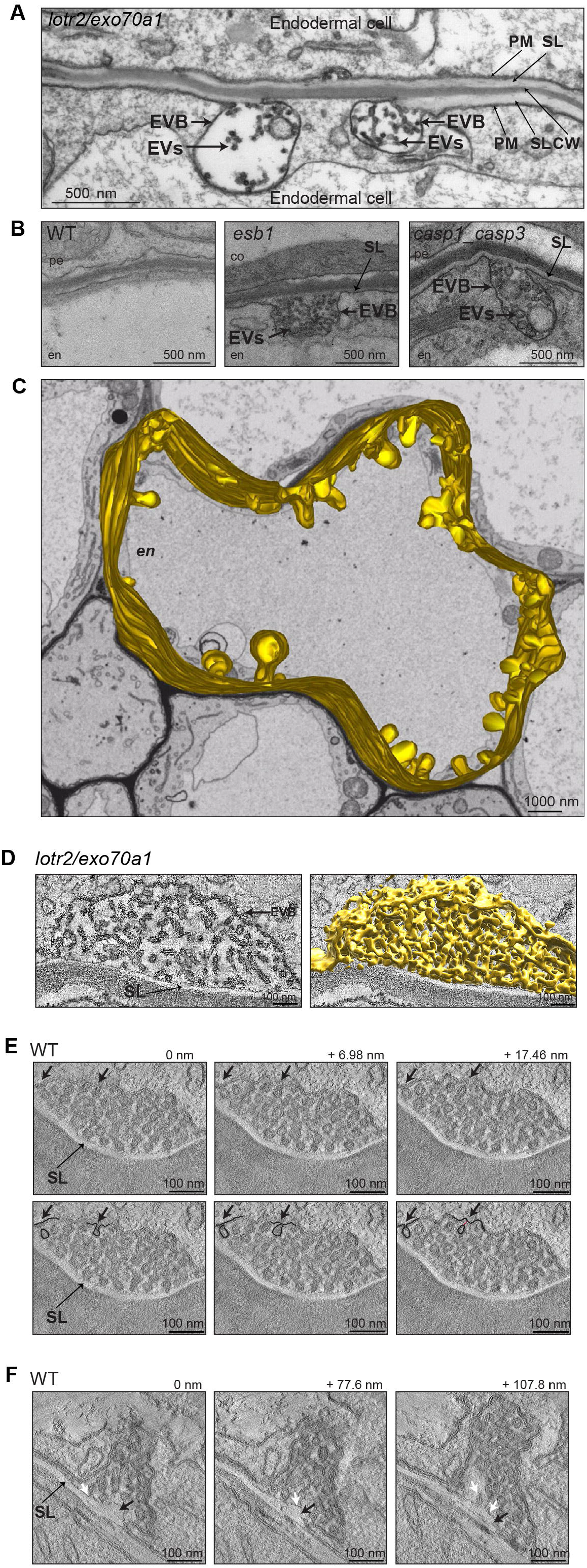
Extracellular vesicular-tubules accumulate in endodermal barriers mutants. **A-D.** TEM sections showing cell wall (CW), suberin lamellae (SL), plasma membrane (PM), extracellular vesicular-tubular (EVs) and EV containing bodies (EVBs) in endodermal (en) cells at 2 mm from root tip. pe, pericycle; co, cortex. A. Endodermal section in lotr2/exo70a1 mutant. B. Endodermal sections in WT, esbl and caspl_casp3 mutants (see also Supplementary Fig. IB). C. 3D model of the PM and its EVBs (highlighted in yellow) in lotr2/exo70a1 mutant. The model was done on a Z portion of 10 Kim starting at 2 mm from the root tip (250 sections, 40 nm thick from a FIB-SEM stack (see also Supplementary Fig. IC,D and Supplementary Movie 1). **D.** Single optical tomography slice and 3D reconstruction of one EVB and its inter-connected vesicular-tubules (segmented in yellow) in lotr2/exo70a1 mutant at 2 mm from tip (see also Supplementary Movie 2). **E.** Series of three optical sections from a tomogram of one EVB in a WT in the suberizing zone. Arrows highlight the invagination of one vesicle (see also Supplementary Movie 3). Lower panels highlight two invaginations events (dark lines) from upper panels (red line highlight a connection between a vesicle and the plasma membrane). **F.** Series of three optical sections from a tomogram of one EVB in a WT in the suberizing zone. Black arrows highlight the growing suberin lamellae, white arrows highlight connections between vesicular-tubules and the suberin lamellae and the small electron dense deposit at the surface of the suberin lamellae that represents the rest of the membrane after fusion (see also Supplementary Movie 4).

The mutants *lotr2/exo70a1, esb1* and *casp1_casp3* all affect CS formation in very different ways. One common feature, however, is that they all display enhanced suberin formation closer to the root tip, where it never occurs in wild-type (Supplementary Fig. 1A) ^20-22^. Indeed, in a number of cases, we could observe a striking association between partially formed suberin lamellae and EVBs (Fig. 1A). We therefore investigated if the accumulation of these vesicles was associated with suberin formation in wild-type. Suberin development is well-described in Arabidopsis roots, with a non-suberized zone (at 2 mm), followed by a patchy zone of ongoing suberization (between 4 and 7 mm), and a fully suberized zone (after 7 mm), where all eight endodermal cells in a section show suberization (Fig. 2A) ^23-25^. Ultrastructural analysis along this developmental gradient revealed a transient, high accumulation of EVBs associated with the endodermal PM in the patchy, suberizing zone (at 5 and 6 mm, Fig. 2B, C), while their number was neglectable prior to suberin formation, as well as in the fully suberized zone. A tight correlation with suberin formation can also be observed in a single root section in the patchy zone, making use of the fact that in this zone, 3 developmental stages of endodermal cells can be simultaneously observed. *i.e.* non-suberized, suberizing and suberized (categorized by the presence, thickness and continuity of suberin lamellae in these cells). Here, we only observed a high number of EVBs in the suberizing cells (Fig. 2D, Supplementary Fig. 2A). The dimensions of the EVBs observed in wild-type, as well as the content and size of their vesicular structures in sections, was similar to the EVBs observed in *lotr2/exo70a1, esb1* and *casp1_casp3* (Supplementary Fig. 2B and C). In order to further strengthen the association between EVBs and suberization, we decided to induce suberin outside of the endodermis by using 1 μM of abscisic acid (ABA) for 14h, which has previously been described to cause suberin accumulation in cortical cells ^24^. Indeed, we observed induction of EVBs in the cortex of ABA-treated plants (Fig. 2E,F, Supplementary Fig. 2D). Again, these EVBs had dimensions, EV content and size matching those observed in the endodermis of CS mutants and suberizing WT endodermal cells (Supplementary Fig. 2B and C). Altogether, our results indicate that suberization is strictly associated with the formation of EVBs.

**Figure 2.**
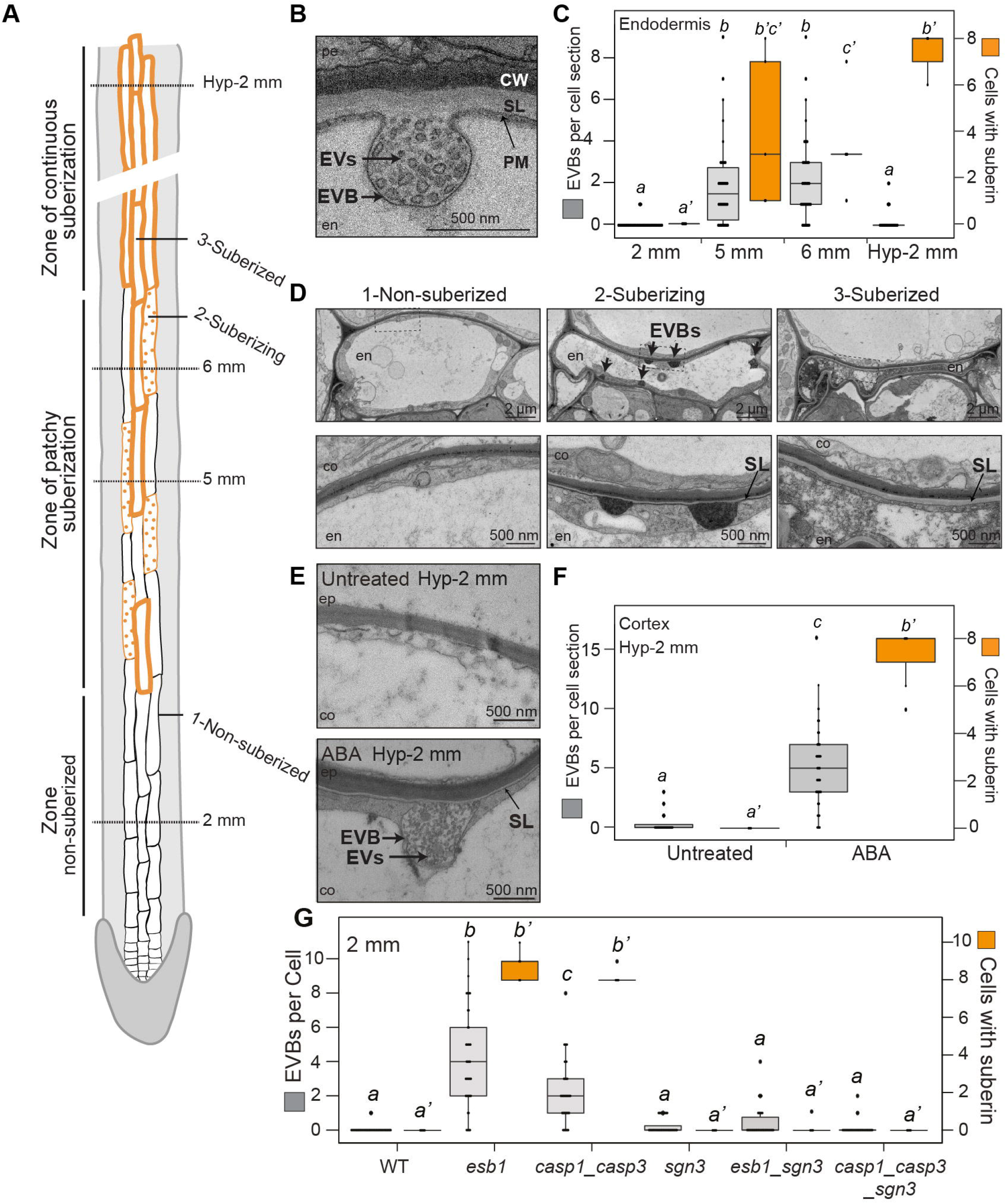
Extracellular vesicular-tubules accumulate in suberizing cells. **A.** Schematic view of suberin differentiation stages in roots. Positions along the roots are marked (2 mm, 5 mm and 6 mm correspond to positions from the root tip and Hyp-2 mm corresponds to position from the hypocotyl-root junction). Examples of non-suberized (1), suberizing (2) and suberized (3) endodermal cells are highlighted. **B,D,E.** TEM sections showing suberin lamellae (SL), EVBs and EVs. en, endodermis; co, cortex; pe, pericycle. B. Endodermal section in WT at 5 mm from root tip. **C,F,G.** Number of visible extracellular vesicular-tubules containing bodies (EVBs), (in grey, left axes) and number of suberized cells (in orange, right axes) in endodermal or cortical layers in full root’s TEM sections. Data are presented as dot plots with box plots overlaid (n?38for EVBs per cells, n?5 for suberized cells per sections). Different letters indicate significant differences between genotypes or growth conditions (P < 0.05). C. Quantifications in the endodermal layer, in WT plants at different positions. D. Pictures illustrating the 3 stages of non-suberized, suberizing and suberized endodermal cells from a root section in the zone of patchy suberization in WT plants (see also Supplementary Fig. 2A). E. Cortical sections in WT plants treated or not with ABA at Hyp-2mm (see also Supplementary Fig. 2D). **F.** Quantifications for the cortical layer at Hyp-2mm in WT plants treated or untreated with ABA. G. Quantifications in the endodermal layer, in WT, esbl and caspl_-casp3, sgn3, esbl _sgn3, and caspl_casp3_sgn3at 2 mm from root tip (data for WT, esbl and caspl-casp3 are also shown in Supplementary Fig. 1 A).

The enhanced suberin deposition of the many CS-defective endodermis mutants is due to stimulation of the CIF/SGN3 (CASPARIAN STRIP INTEGRITY FACTOR / SCHENGEN3) pathway^22,26,27^. Consequently, the *sgn3* mutant does not display enhanced suberin formation and is epistatic to *esb1* and *casp1_casp3* (Fig. 2G, Supplementary Fig. 3A). We therefore tested whether, EVB formation in the early differentiating endodermis of CS mutants was also suppressed in *sgn3_esb1* and *sgn3_casp1_casp3* mutants. Indeed, neither *sgn3* nor *sgn3_esb1* and *sgn3_casp1_casp3* mutants displayed EVBs in the early differentiated endodermis (Fig. 2G, Supplementary Fig. 3A). Thus, the strict correlation between EVB presence and suberin formation holds up even when challenged by a second stimulation of suberization, independent from ABA ^28^ - in this case, peptide receptor-mediated. Together, our data is strongly suggesting a causal relationship between EVBs and suberin deposition.

Suberin is a polyester that is formed as a secondary cell wall, deposited in the form of lamellae just outside of the plasma membrane. Its monomeric precursors are thought to be produced at the endoplasmic reticulum and to be polymerized in the apoplast ^18,29^. However, the transport of hydrophobic suberin monomers to the apoplast is poorly understood. Current research focusses mainly on the role of ATP-binding cassette (ABC) transporters and lipid transfer proteins (LTPs), but their functional significance for suberin deposition remains to be demonstrated ^18,30^. Alternatively, a key role of vesicle-mediated secretion for suberin export has repeatedly been hypothesized, taking into account their hydrophobic nature ^18,29^. However, little evidence has been provided for this and earlier observations of EVB-like vesicles in differentiating endodermal and exodermal cells ^31,32^, have never entered the literature as evidence for secretion-based suberization ^18,29^. We therefore wondered if the vesicles accumulating in suberizing cells reflect suberin monomer secretion to the apoplast. The lipidic nature of suberin allows its staining in whole-mount roots with Fluorol yellow ^23-25^. A close look at Fluorol yellow staining in the first suberizing cells of untreated or ABA-treated roots showed a signal not only at the cell periphery (apoplast) but also as punctate structures (Fig. 3A). These structures are smaller than 1 μm and could therefore correspond to the large vesicles observed at the ultrastructural level in suberizing cells. However, Fluorol yellow staining uses harsh conditions and observed subcellular structures cannot be straightforwardly compared with live cell structures or combined with ultrastructural analysis. Nevertheless, since apoplastic suberin, even in the early stages of suberization, occurs exclusively in lamellae, we reasoned that the presence of punctate structures, stained by Fluorol yellow in early suberizing cells, lends at least some support to the notion of a vesicular, lipidic cargo intermediate during suberization.

**Figure 3.**
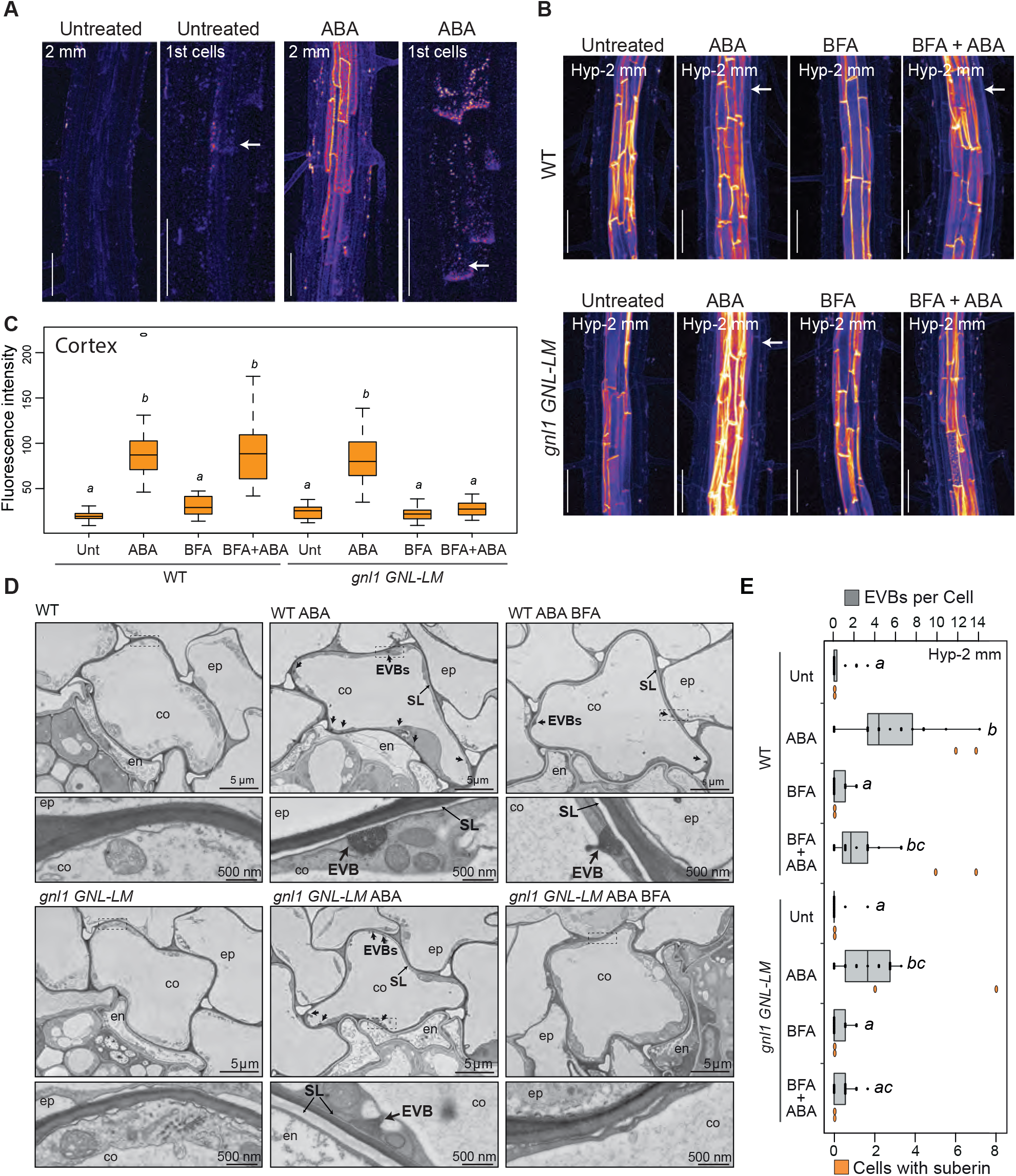
Secretion-dependent suberin deposition. **A-B.** Fluorol yellow staining for suberin in roots. Fluorescence is presented as LUT (Fire), scale bars, 50 μm. **A.** WT plants treated or not with ABA. For each conditions, left pictures taken at 2 mm from root tip, right pictures display the first suberizing cells (1st cells, highlighted with arrows). **B-C.** WT and gnl1GNL-LM lines treated or not with ABA and/or BFA. **B.** Pictures taken at Hyp-2mm from hypocotyl. Arrows highlight the cortical suberin. **C.** Quantification of maximum fluorescence intensity in cortical-epidermal walls, data presented as box plots (n≥10), different letters indicate significant differences between genotypes or growth conditions (P < 0.05). **D.** TEM sections showing a cortical cell in WT and gnl1GNL-LM lines treated or not with ABA and ABA+BFA at Hyp-2mm. Arrows highlight EVBs. Lower panels correspond to a magnification from upper panels (zone defined with dashed lines). **E.** Number of EVBs (in grey, upper axes) and number of suberized cells (in orange, lower axes) in cortical layers in full root’s TEM sections in WT and gnl1GNL-LM lines treated or not with ABA and/or BFA. Data are presented as dot plots with box plots overlaid (n≥16 endodermal cells for EVBs per endodermal cells, n=2 for suberized cells per sections). Different letters indicate significant differences between genotypes and growth conditions for EVB (P < 0.05).

In order to study the role of secretory endomembrane trafficking in suberin deposition, we thought to make use of the fact that ABA induces *de novo* suberin formation in cortical cells ^24^, allowing us to compare the same cell type in an induced and un-induced state. To address the origin and identity of the EVs, we screened the Wave Line collection of subcellular markers ^33^ upon ABA treatment, but failed to observe cortex-specific changes in fluorescence in any of these lines after treatment. This was surprising, since the marker collection covers the major intracellular membrane compartments, such as Golgi, TGN, recycling endosomes, vacuoles and MVBs (Supplementary Fig. 3B). We then attempted to affect endomembrane trafficking and secretion and study its consequences on suberization. We refrained from using constitutive trafficking mutants, since they are either weak or have such severe pleiotropic defects that they are not able to undergo proper embryogenesis and root development (e.g. ^34-36^). Since suberization is highly responsive to many different stresses, we predicted that it would be impossible to separate primary from secondary effects in these mutants. We therefore focused on trafficking inhibitors in order to allow for a more acute manipulation of membrane trafficking. Brefeldin A (BFA), is a well-characterized inhibitor of membrane trafficking whose mechanism of action on GDP/GTP exchange factors (GEFs) for ARF G-proteins are understood. Importantly, single point mutations can predictably render different ARF-GEFs-acting at different points of the trafficking pathway - either resistant or sensitive to BFA. This effectively allows to use the same inhibitor to be largely specific to endosomal trafficking in WT or to affect trafficking already at ER-to-cis-Golgi trafficking, depending on the genetic background used ^37-39^. We first used BFA on WT, in combination with ABA treatment, allowing us to observe, *de novo* suberin formation in cortical cells. In WT, BFA treatment did not affect ABA-induced cortical suberization, nor did it decrease the quantity of EVB structures (Fig. 3B-E). However, in *gnl1-1GNL1-LM* (*gnom-like1*) plants, a genotype with root development indistinguishable from wild-type ^38^, ABA-dependent suberin deposition in cortical cell walls was blocked upon BFA treatment (Fig. 3B,C). Importantly, BFA treatment also abrogated the increase of EVBs in cortical cells, after ABA treatment in *gnl1-1GNL1-LM* (Fig. 3 D,E). *gnl1-1GNL1-LM* is hypersensitive to Brefeldin A (BFA), blocking secretory trafficking already at the ER-to-Golgi step ^38,40^. Thus, early secretory endomembrane trafficking is required for suberin deposition in the cell wall, whereas affecting exclusively endosomal trafficking appears to be ineffective to block EVB formation and suberin deposition.

It has been proposed previously for the process of EVB biogenesis that these structures might stem from the MVB/LE pathway ^1,6^. Support for this model comes from the overall structural and topological resemblance of EVBs and MVBs. However, we were unable to observe any comparable presence of MVBs, as we see for EVBs in suberizing cells. Moreover, the massive increase in EVBs upon ABA treatment in cortical cells is not reflected in observable changes in MVB/LE numbers or size, when using our Wave marker lines (Supplementary Fig. 3B). Lastly, interfering with endosomal trafficking by BFA treatment in wild-type, did not affect EVB formation or suberization (Fig 3 B-D). By contrast, inhibiting early secretory trafficking at the level of the ER by BFA treatment of the *gnl1 GNL1-LM* mutant did affect EVB formation and suberization (Fig. 3-E). Moreover, direct transfer of lipids between ER and PM is known to occur at ER-PM contact sites and to be crucial for cellular membrane homeostasis^41,42^. It is therefore straightforward to assume that the lipid-like suberin precursors, which are synthesized at the ER, are transferred directly between ER and PM. Similar to lipid droplets, shown to burgeon from the ER ^43^, luminal tubules rich in suberin monomer lipids might initially form at their site of biogenesis in the ER and then fuse with the PM, placing suberin monomer-containing tubules into the apoplast where they can be polymerized (Fig. 4). Intriguingly, lipid droplet formation also involves GPATs (GLYCEROL-3-PHOSPHATE SN-2-ACYLTRANSFERASES), the same enzyme class that is catalyzing the last step(s) of suberin/cutin monomer biosynthesis ^44^. In addition, lipid droplet homeostasis was shown to depend on the ARF1/COPI machinery ^44^ which is precisely the step sensitized to BFA action in the *gnl1 /GNL1-LM* background. Such a model of extracellular tubule-mediated monomer transport draws tantalizing parallels to the extracellular tubulo-vesicular structures observed in arbuscules at the fungal plant membrane interface ^8,9^. There is good evidence that lipid-like molecules are provided to the fatty acid-auxotrophic fungus and that their generation requires GPAT enzymes ^45^. We therefore speculate that there might be a deep cellular and molecular resemblance of the mechanisms providing lipidic molecules to the fungus at the arbuscule and the process of transporting very similar, lipid-like suberin monomers to the apoplast during suberization. Our work here establishes a developmental role for EVBs in the formation of a major cell wall polymer, distinct from the currently reported functions of EVBs in direct, symbiotic and pathogenic interactions.

**Figure 4.**
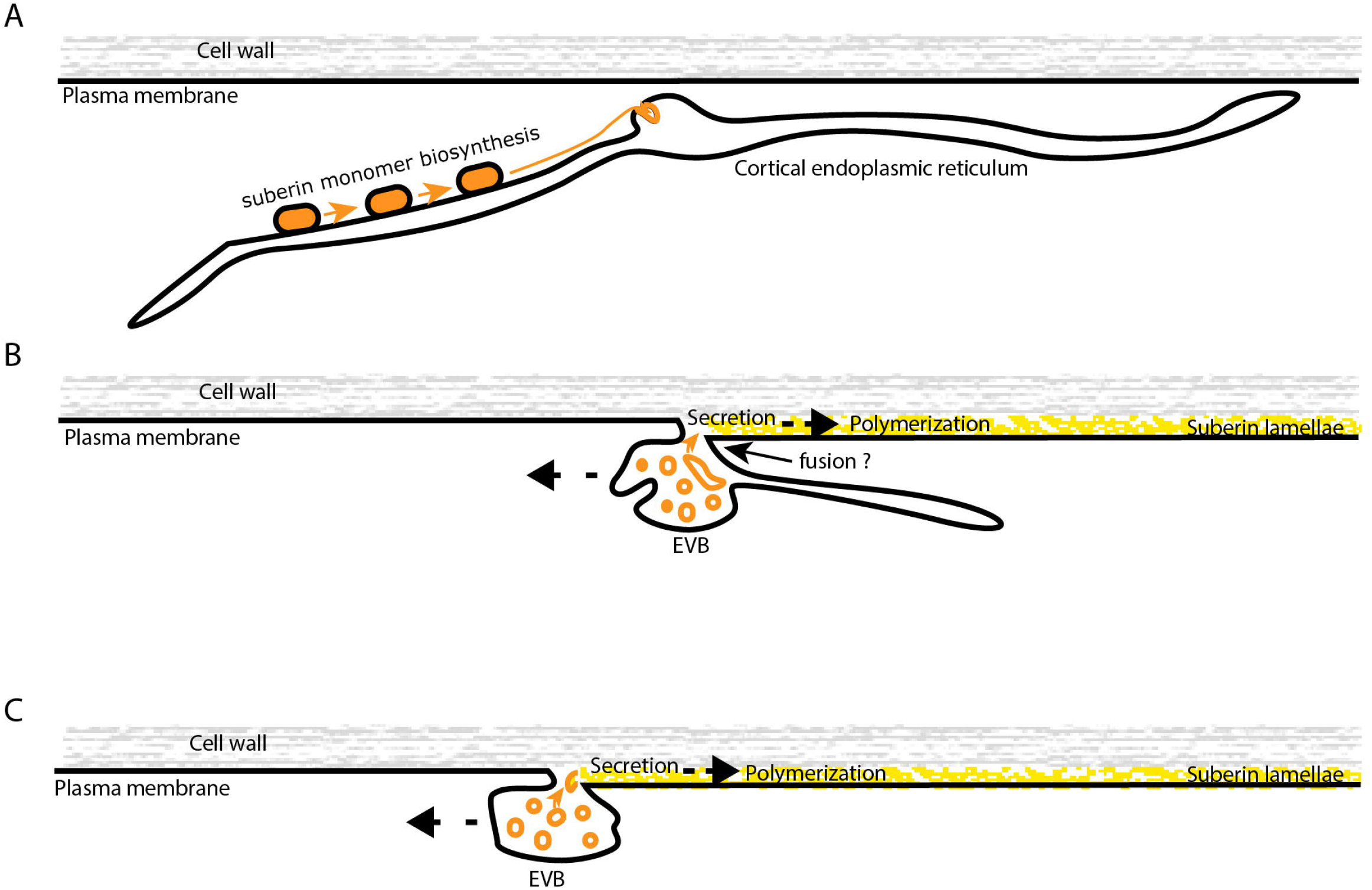
Speculative model for suberin monomer transport by vesiculo-tubular intermediates. **A.** The lipid-like suberin monomers produced by the successive activities of their biosynthetic enzymes (orange blocks) at the endoplasmic reticulum (ER) might associate into cortical ER-derived subdomains (orange) that evaginate into the lumen of the ER. B. The initially ER-derived structure swells, accommodating larger amounts of monomer-containing tubules (orange) and eventually disconnects from the ER and fuses with the nearby plasma membrane. C The suberin monomer containing tubules are placed in the apoplast and are gradually consumed as substrates of cell wall localized suberin polymerizing enzymes, forming suberin lamellae (yellow).

## Acknowledgments

We thank the Central Imaging Facility (CIF) for expert technical support. Caroline Kizilyaprak is thanked for her crucial help during FIB image acquisition, Willy Blanchard and Christel Genoud for their help in modeling. We are thankful to, Gerd Jürgens for sharing published material. This work was supported by Société Académique Vaudoise to L.K., and SNF (grant 310030B_176399) to N.G., (grants 31003A_179159 and PCEGP3_187007) to M.B.

## Author contributions

M.B and N.G designed the project, DDB performed EM analysis, LK and JD performed initial work on *exo70a1/lotr2* characterization, PM, analyzed endomembrane markers, MB, prepared material, performed suberin analysis, analyzed data with DDB and wrote the manuscript with NG.

## Experimental Procedures

### Material

All experiments were performed with Arabidopsis, ecotype Columbia. Mutants and transgenic lines analyzed in this study were generated and characterized before: *exo70a1-4/lotr2-1* ^20^, *sgn3-3* ^22^, *esb1-1* ^21^, *casp1_casp3* ^46^, *sgn3-3_esb1-1* ^22^, *sgn3-3_casp1_casp3* ^22^, *gnl1-1GNL1-LM* ^38^, and the following Wave lines ^33^ Wave2Y (*UBQ10::RabF2b/ARA7-EYFP*), Wave3Y (*UBQ10::RabC1-EYFP*), Wave5Y (*UBQ10::RabG3f-EYFP*), Wave7Y (*UBQ10::RabF2a/Rha1-EYFP*) and Wave13Y (*UBQ10::VTI12-EYFP*). The corresponding gene numbers are as follows: *CASP1*, At2g36100; *CASP3*, At2g27370; *ESB1*, At2g28670; *EXO70A1/LOTR2*, At5g03540; *GNL1*, At5g39500; *GPAT5*, At3g11430; *SGN3*, At4g20140.

### Growth conditions

For all experiments, seeds were surface sterilized, sown on 0.5 x MS (Murashige and Skoog) 0.8% agar plates, incubated 2 to 3 days at 4°C and grown vertically in growth chambers at 22°C, under continuous light (100 μE). All analyses were performed on 5-day-old seedlings. Treatments were performed as transfers for 16 h in a way that seedlings were 5-day-old at the point of analysis. ABA and BFA were directly added to 0.5 x MS plates at the following concentrations: 1 μM and 25 μM, respectively, for 14 h.

### TEM analysis

Plants were fixed in glutaraldehyde solution (EMS, Hatfield, PA) 2.5% in phosphate buffer (PB 0.1 M [pH 7.4]) for 1 h at RT and postfixed in a fresh mixture of osmium tetroxide 1% (EMS, Hatfield, PA) with 1.5% of potassium ferrocyanide (Sigma, St. Louis, MO) in PB buffer for 1 h at RT. The samples were then washed twice in distilled water and dehydrated in ethanol solution (Sigma, St Louis, MO, US) at graded concentrations (30% - 40 min; 50% - 40 min; 70% - 40 min; 100% - 2x 1 h). This was followed by infiltration in Spurr resin (EMS, Hatfield, PA, US) at graded concentrations (Spurr 33% in ethanol - 4 h; Spurr 66% in ethanol - 4 h; Spurr 100% - 2x 8 h) and finally polymerized for 48 h at 60°C in an oven. Ultrathin sections of 50 nm thickness were cut transversally at 2, 5, and 6 mm from the root tip and at 2 mm below the hypocotyl-root junction, using a Leica Ultracut (Leica Mikrosysteme GmbH, Vienna, Austria), picked up on a copper slot grid 2×1 mm (EMS, Hatfield, PA, US) coated with a polystyrene film (Sigma, St Louis, MO, US). Sections were post-stained with uranyl acetate (Sigma, St Louis, MO, US) 4% in H_2_O for 10 min, rinsed several times with H_2_O followed by Reynolds lead citrate in H_2_O (Sigma, St Louis, MO, US) for 10 min and rinsed several times with H_2_O. Micrographs were taken with a transmission electron microscope Philips CM100 (Thermo Fisher Scientific, Waltham, MA USA) at an acceleration voltage of 80 kV with a TVIPS TemCamF416 digital camera (TVIPS GmbH, Gauting, Germany) using the software EM-MENU 4.0 (TVIPS GmbH, Gauting, Germany). Panoramic alignment were performed with the software IMOD ^47^.

### Focused Ion Beam Scanning Electron Microscopy (FIB-SEM)

The resin block was oriented and mounted on an aluminium support, glued with conductive resin Epotek H2OS^®^ (EMS, Hatfield, PA, US), and polymerized overnight in an oven at 60°C. It was then trimmed in the ultramicrotome to position the sample (2mm from the root tip) and prepare its geometry for FIB-SEM analysis. 30 nm of platinum was then sputter coated on the block using a Leica EM SCD 500 sputter coater (Leica Mikrosysteme GmbH, *Vienna*, Austria). Serial block face imaging is finally performed in a Helios NanoLab 650 (Thermo Fisher Scientific, Waltham, MA USA), using the FEI Slice and View software™. The milling of 40 nm slice thickness was done at 30 kV acceleration voltage and 6.6 nA current. The cross section images were acquired by detecting backscattered electrons with the In-column detector (ICD) in immersion mode, at 4.2 mm of working distance and an electron beam of 2 kV, 800 pA and 5 μs of dwell time with a frame of 4096 × 3536 pixels, an horizontal field width of 56 μm and a pixel size of 13.6 nm, total Z volume acquired is 27.96 μm. Further details on block geometry and milling strategy were previously described in ^48^. Volume alignment and 3D modelling were performed using IMOD software ^47^.

### TEM tomography and 3D reconstruction

For electron tomography, semi-thin sections of 250nm thickness were cut transversally to the root using a Leica Ultracut (Leica Mikrosysteme GmbH, Vienna, Austria) and then, picked up on 75 square mesh copper grids (EMS, Hatfield, PA, US). Sections were post-stained on both sides with uranyl acetate (Sigma, St Louis, MO, US) 2% in H_2_O for 10 min and rinsed several times with H_2_O. Protein A Gold 10nm beads (Aurion, Wageningen, The Netherlands) were applied as fiducials on both sides of the sections and the grids were placed on a dual axis tomography holder (Model 2040, Fischione Instruments). The aera of interest was taken with a transmission electron microscope JEOL JEM-2100Plus (JEOL Ltd., Akishima, Tokyo, Japan) at an acceleration voltage of 200 kV with a TVIPS TemCamXF416 digital camera (TVIPS GmbH, Gauting, Germany) using the SerialEM software package^49^. Micrographs were taken as single or dual-axis tilt series over a range of −60° to +60° using SerialEM at tilt angle increment of 1°. Tomogram reconstruction was done with IMOD software ^47^, segmentation with Ilastik software package ^50^ and model visualization with Imaris software package (Oxford Instruments).

### Fluorescence microscopy

Fluorol yellow (FY) staining was used to visualize suberin in whole-mounted roots as described before ^25^. Seedlings were incubated in freshly prepared FY 088 (0.01% w/v, lactic acid) at 70°C for 30 min, rinsed with water and counterstained with aniline blue (0.5% w/v, water) at RT for 30 min in darkness, washed, mounted in 70% glycerol and observed with confocal. For visualization of cell files in live imaging, 5-day-old seedlings were incubated in the dark for 10 min in a fresh solution of 15 mM (10 mg/ml) Propidium Iodide (PI) dissolved in liquid 0.5 x MS and rinsed in liquid 0.5 x MS prior to imaging. Confocal laser scanning microscopy experiments were performed on a Zeiss LSM 700 or Zeiss LSM 880. Excitation and detection windows were set as follows: FY 488 nm, SP 640 nm; YFP 488 nm, 500-530 nm; PI 555 nm, SP 640 nm. For FY imaging laser power was reduced as low as 0.2% to limit bleaching. Confocal pictures were subsequently analyzed with Fiji ^51^, channels merged, Z stacks converted as 3D-projections and/or orthogonal views. FY staining is presented as Look Up Tables (LUT). Fluorescence in the cortex was quantified from pictures taken with exactly the same parameters, by tracing a 10 μm line crossing the cell wall between epidermal and cortical cells, measuring the fluorescence intensity and considering the maximum value per measure.

### Statistical analysis

All graphics and statistical analyses were done in the R environment. For multiple comparisons between genotypes or conditions, one-way ANOVA was performed, and Tukey’s test subsequently used as a multiple comparison procedure. When the data did not follow the linear model assumption Kruskal-Wallis and nonparametric Tukey’s test were performed for multiple comparison.

## Supplementary movie legends

**Supplementary Movie 1.**

**3D reconstruction of the PM and its EVBs in *lotr2/exo70a1* mutant.** Travelling in a series of 40 nm FIB-SEM slices through a Z volume of 10 μm showing the high number of extracellular vesicular-tubules containing bodies (EVBs) in an endodermal suberizing cell of a *lotr2/exo70a1* mutant at 2 mm from the root tip. The 3D model in yellow highlight the PM and its EVBs. The overview picture is shown in the Fig. 1C, some screen shots in Supplementary Fig. 1C and orthogonal views in Supplementary Fig. 1D.

**Supplementary Movie 2.**

**3D reconstruction of one EVB in *lotr2/exo70a1* mutant.** Travelling in a series of 0.77 nm optical tomography slices through a Z volume of 191 nm showing the high number of inter-connected vesicular-tubules inside one EVB in an endodermal suberizing cell of a *lotr2/exo70a1* mutant at 2 mm from the root tip. The 3D model in yellow highlight one EVB. A single optical tomography slice and an overview picture of the model is shown in the Fig.1D.

**Supplementary Movie 3.**

**3D reconstruction of one EVB in WT.** Travelling in a series of 0.38 nm optical tomography slices through a Z volume of 162 nm showing one EVB in an endodermal suberizing cell from a WT root in the suberizing zone. Serie of three optical sections is shown in Fig.1E.

**Supplementary Movie 4.**

**3D reconstruction of one EVB in WT.** Travelling in a series of 0.38 nm optical tomography slices through a Z volume of 148 nm showing one EVB and the growing suberin lamellae in an endodermal suberizing cell from a WT root in the suberizing zone. Serie of three optical sections is shown in Fig.1F.

## References

1. van Niel, G., D’Angelo, G. & Raposo, G. Shedding light on the cell biology of extracellular vesicles. Nature Reviews Molecular Cell Biology 19, 213–228 (2018).

2. Herman, E. M. & Lamb, C. J. Arabinogalactan-Rich Glycoproteins Are Localized on the Cell Surface and in Intravacuolar Multivesicular Bodies. Plant Physiology 98, 264–272 (1992).

3. Marchant, R. & Robards, A. W. Membrane Systems Associated with the Plasmalemma of Plant Cells. Annals of Botany 32, 457–471 (1968).

4. Esau, K., Cheadle, V. I. & Gill, R. H. Cytology of Differentiating Tracheary Elements li. Structures Associated with Cell Surfaces. American Journal of Botany 53, 765–771 (1966).

5. Baldrich, P. et al. Plant Extracellular Vesicles Contain Diverse Small RNA Species and Are Enriched in 10-to 17-Nucleotide “Tiny” RNAs. The Plant Cell 31, 315–324 (2019).

6. Cai, Q., He, B., Weiberg, A., Buck, A. H. & Jin, H. Small RNAs and extracellular vesicles: New mechanisms of cross-species communication and innovative tools for disease control. PLOS Pathogens 15, e1008090 (2019).

7. Cai, Q. et al. Plants send small RNAs in extracellular vesicles to fungal pathogen to silence virulence genes. Science 360, 1126–1129 (2018).

8. Ivanov, S., Austin, J., Berg, R. H. & Harrison, M. J. Extensive membrane systems at the host-arbuscular mycorrhizal fungus interface. Nature Plants 5, 194–203 (2019).

9. Roth, R. et al. Arbuscular cell invasion coincides with extracellular vesicles and membrane tubules. Nature Plants 5, 204–211 (2019).

10. Meyer, D., Pajonk, S., Micali, C., O’Connell, R. & Schulze-Lefert, P. Extracellular transport and integration of plant secretory proteins into pathogen-induced cell wall compartments. The Plant Journal 57, 986–999 (2009).

11. Rutter, B. D. & Innes, R. W. Extracellular Vesicles Isolated from the Leaf Apoplast Carry Stress-Response Proteins. Plant Physiology 173, 728–741 (2017).

12. An, Q., Hückelhoven, R., Kogel, K.-H. & Bel, A. J. E. V. Multivesicular bodies participate in a cell wall-associated defence response in barley leaves attacked by the pathogenic powdery mildew fungus. Cellular Microbiology 8, 1009–1019 (2006).

13. Regente, M. et al. Plant extracellular vesicles are incorporated by a fungal pathogen and inhibit its growth. Journal of Experimental Botany 68, 5485–5495 (2017).

14. de la Canal, L. & Pinedo, M. Extracellular vesicles: a missing component in plant cell wall remodeling. Journal of Experimental Botany 69, 4655–4658 (2018).

15. Kim, S.-J. & Brandizzi, F. The plant secretory pathway: an essential factory for building plant cell walls. Plant Cell Physiol pct197 (2014) doi:10.1093/pcp/pct197.

16. Liljegren, S. J. et al. Regulation of membrane trafficking and organ separation by the NEVERSHED ARF-GAP protein. Development 136, 1909–1918 (2009).

17. Goh, T. et al. VPS9a, the Common Activator for Two Distinct Types of Rab5 GTPases, Is Essential for the Development of *Arabidopsis thaliana*. The Plant Cell 19, 3504–3515 (2007).

18. Vishwanath, S. J., Delude, C., Domergue, F. & Rowland, O. Suberin: biosynthesis, regulation, and polymer assembly of a protective extracellular barrier. Plant Cell Reports 34, 573–586 (2015).

19. Alassimone, J. et al. Polarly localized kinase SGN1 is required for Casparian strip integrity and positioning. Nat Plants 2, 16113 (2016).

20. Kalmbach, L. et al. Transient cell-specific EXO70A1 activity in the CASP domain and Casparian strip localization. Nature Plants 3, 17058 (2017).

21. Hosmani, P. S. et al. Dirigent domain-containing protein is part of the machinery required for formation of the lignin-based Casparian strip in the root. Proc. Natl. Acad. Sci. U.S.A. 110, 14498–14503 (2013).

22. Pfister, A. et al. A receptor-like kinase mutant with absent endodermal diffusion barrier displays selective nutrient homeostasis defects. Elife 3, e03115 (2014).

23. Andersen, T. G. et al. Diffusible repression of cytokinin signalling produces endodermal symmetry and passage cells. Nature 555, 529–533 (2018).

24. Barberon, M. et al. Adaptation of Root Function by Nutrient-Induced Plasticity of Endodermal Differentiation. Cell 164, 447–459 (2016).

25. Naseer, S. et al. Casparian strip diffusion barrier in Arabidopsis is made of a lignin polymer without suberin. Proc. Natl. Acad. Sci. U.S.A. 109, 10101–10106 (2012).

26. Doblas, V. G. et al. Root diffusion barrier control by a vasculature-derived peptide binding to the SGN3 receptor. Science 355, 280–284 (2017).

27. Fujita, S. et al. SCHENGEN receptor module drives localized ROS production and lignification in plant roots. The EMBO Journal 18 (2020).

28. Shukla, V. et al. Suberin plasticity to developmental and exogenous cues is regulated by a set of MYB transcription factors. bioRxiv 2021.01.27.428267 (2021) doi:10.1101/2021.01.27.428267.

29. Franke, R. & Schreiber, L. Suberin — a biopolyester forming apoplastic plant interfaces. Current Opinion in Plant Biology 10, 252–259 (2007).

30. Yadav, V. et al. ABCG Transporters Are Required for Suberin and Pollen Wall Extracellular Barriers in Arabidopsis. The Plant Cell 26, 3569–3588 (2014).

31. Ma, F. & Peterson, C. A. Development of cell wall modifications in the endodermis and exodermis of *Allium cepa* roots. Canadian Journal of Botany 79, 621–634 (2001).

32. Scott, M. G. & Peterson, R. L. The root endodermis in *Ranunculus acrís*. I. Structure and ontogeny. Canadian Journal of Botany 57, 1040–1062 (1979).

33. Geldner, N. et al. Rapid, combinatorial analysis of membrane compartments in intact plants with a multicolor marker set. Plant J. 59, 169–178 (2009).

34. Roudier, F. et al. Very-long-chain fatty acids are involved in polar auxin transport and developmental patterning in Arabidopsis. Plant Cell 22, 364–375 (2010).

35. Jaillais, Y., Fobis-Loisy, I., Miège, C., Rollin, C. & Gaude, T. AtSNX1 defines an endosome for auxin-carrier trafficking in Arabidopsis. Nature 443, 106–109 (2006).

36. Geldner, N. et al. Partial loss-of-function alleles reveal a role for GNOM in auxin transport-related, post-embryonic development of Arabidopsis. Development 131, 389–400 (2004).

37. Richter, S. et al. Polarized cell growth in Arabidopsis requires endosomal recycling mediated by GBF1-related ARF exchange factors. Nat. Cell Biol. 14, 80–86 (2012).

38. Richter, S. et al. Functional diversification of closely related ARF-GEFs in protein secretion and recycling. Nature 448, 488–492 (2007).

39. Geldner, N. et al. The Arabidopsis GNOM ARF-GEF mediates endosomal recycling, auxin transport, and auxin-dependent plant growth. Cell 112, 219–230 (2003).

40. Teh, O.-K. & Moore, I. An ARF-GEF acting at the Golgi and in selective endocytosis in polarized plant cells. Nature 448, 493–496 (2007).

41. Wu, H., Carvalho, P. & Voeltz, G. K. Here, there, and everywhere: The importance of ER membrane contact sites. Science 361, eaan5835 (2018).

42. Stefan, C. J., Manford, A. G. & Emr, S. D. ER-PM connections: sites of information transfer and inter-organelle communication. Current Opinion in Cell Biology 25, 434–442 (2013).

43. Walther, T. C., Chung, J. & Jr, R. V. F. Lipid Droplet Biogenesis. 22 (2017).

44. Wilfling, F. et al. Arf1/COPI machinery acts directly on lipid droplets and enables their connection to the ER for protein targeting. eLife 3, e01607 (2014).

45. Luginbuehl, L. H. et al. Fatty acids in arbuscular mycorrhizal fungi are synthesized by the host plant. Science 356, 1175–1178 (2017).

46. Roppolo, D. et al. A novel protein family mediates Casparian strip formation in the endodermis. Nature 473, 380–383 (2011).

47. Kremer, J. R., Mastronarde, D. N. & McIntosh, J. R. Computer Visualization of Three-Dimensional Image Data Using IMOD. Journal of Structural Biology 116, 71–76 (1996).

48. Kizilyaprak, C., Longo, G., Daraspe, J. & Humbel, B. M. Investigation of resins suitable for the preparation of biological sample for 3-D electron microscopy. Journal of Structural Biology 189, 135–146 (2015).

49. Mastronarde, D. N. Automated electron microscope tomography using robust prediction of specimen movements. Journal of Structural Biology 152, 36–51 (2005).

50. Berg, S. et al. ilastik: interactive machine learning for (bio)image analysis. Nature Methods 16, 1226–1232 (2019).

51. Schindelin, J. et al. Fiji: an open-source platform for biological-image analysis. Nature Methods 9, 676–682 (2012).

