## Supplementary figures and images for "Extracellular membrane tubules involved in suberin deposition in plant cell walls"

### Supplementary Fig. 1

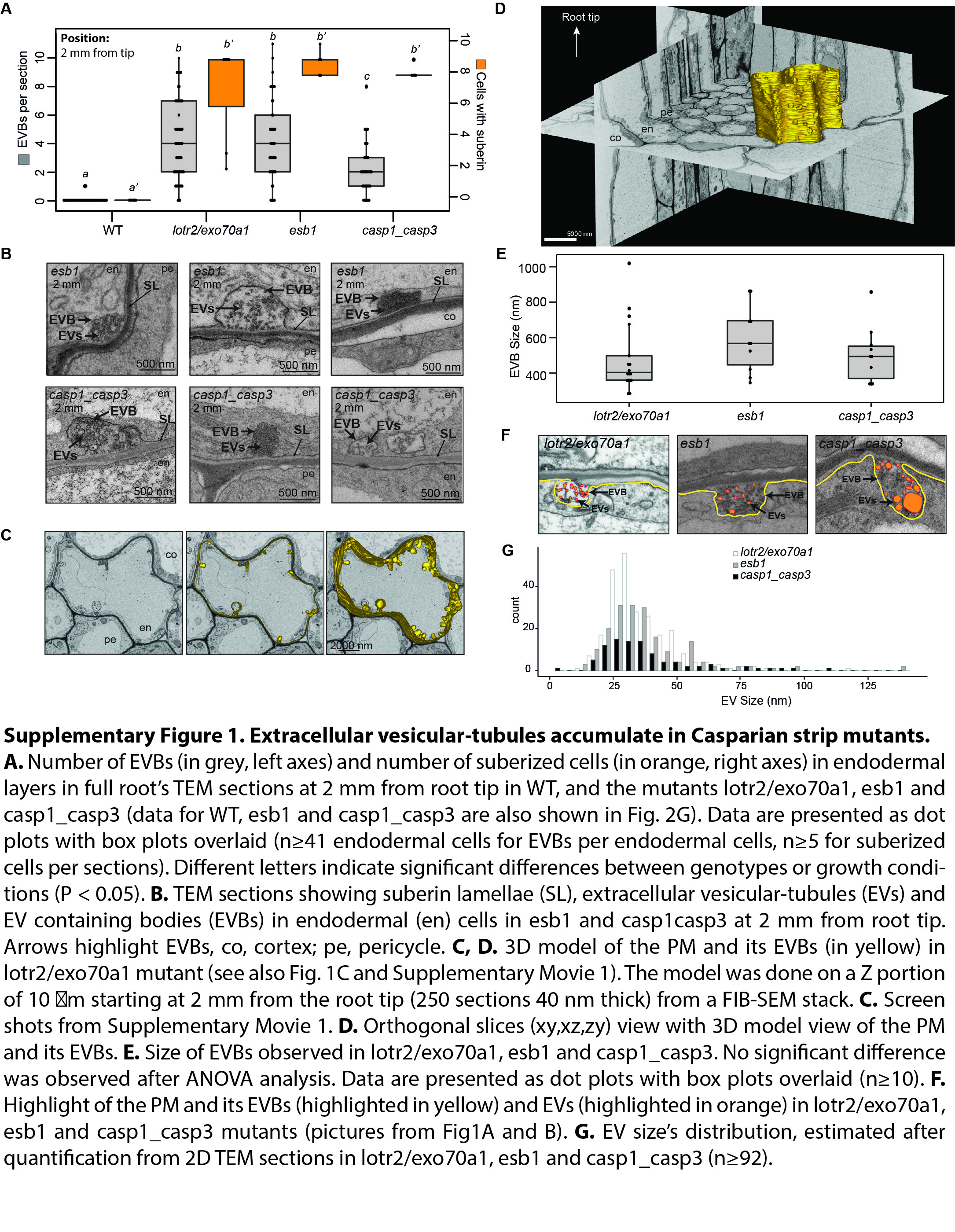

### Supplementary Fig. 2

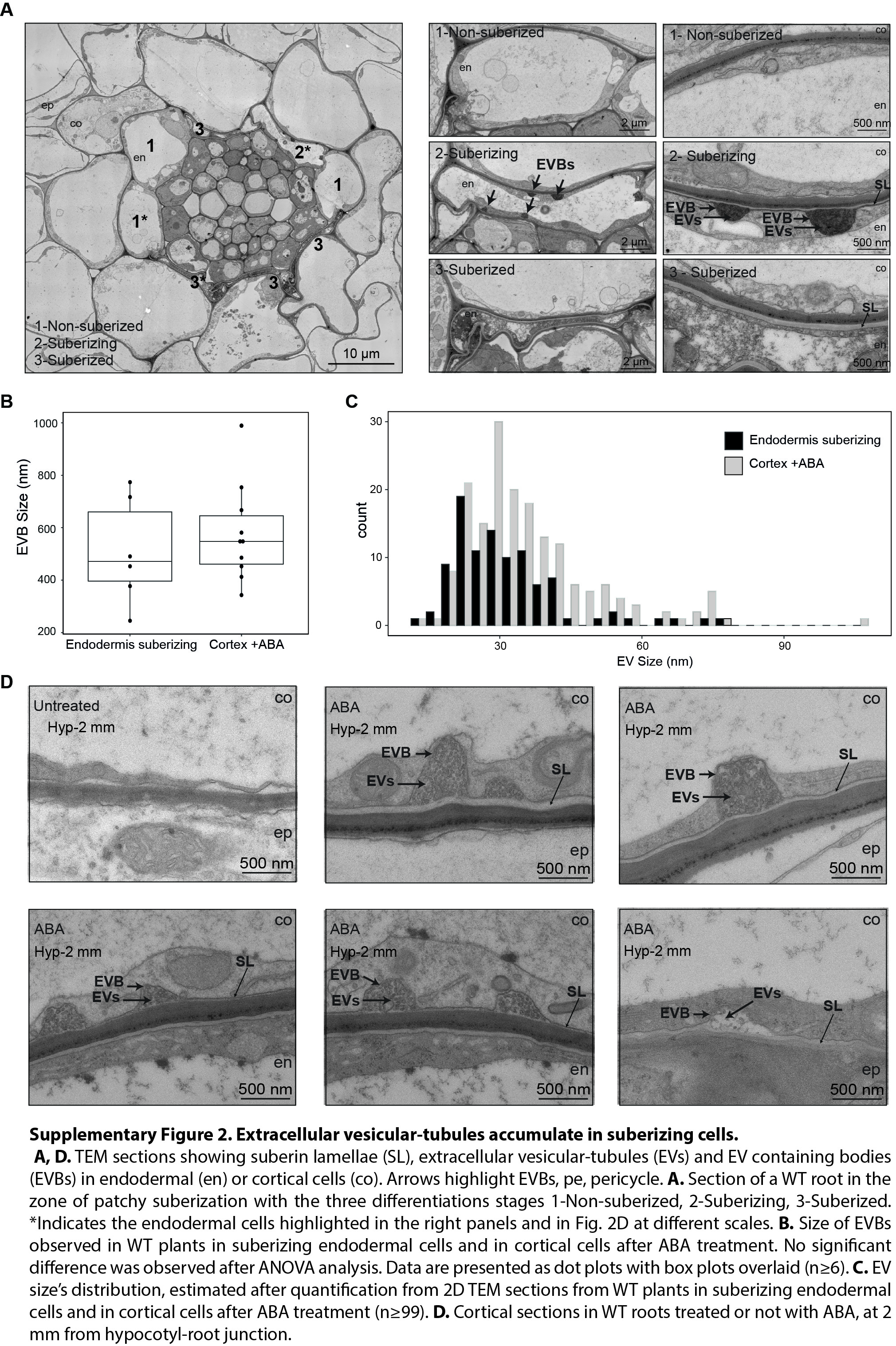

### Supplementary Fig. 3

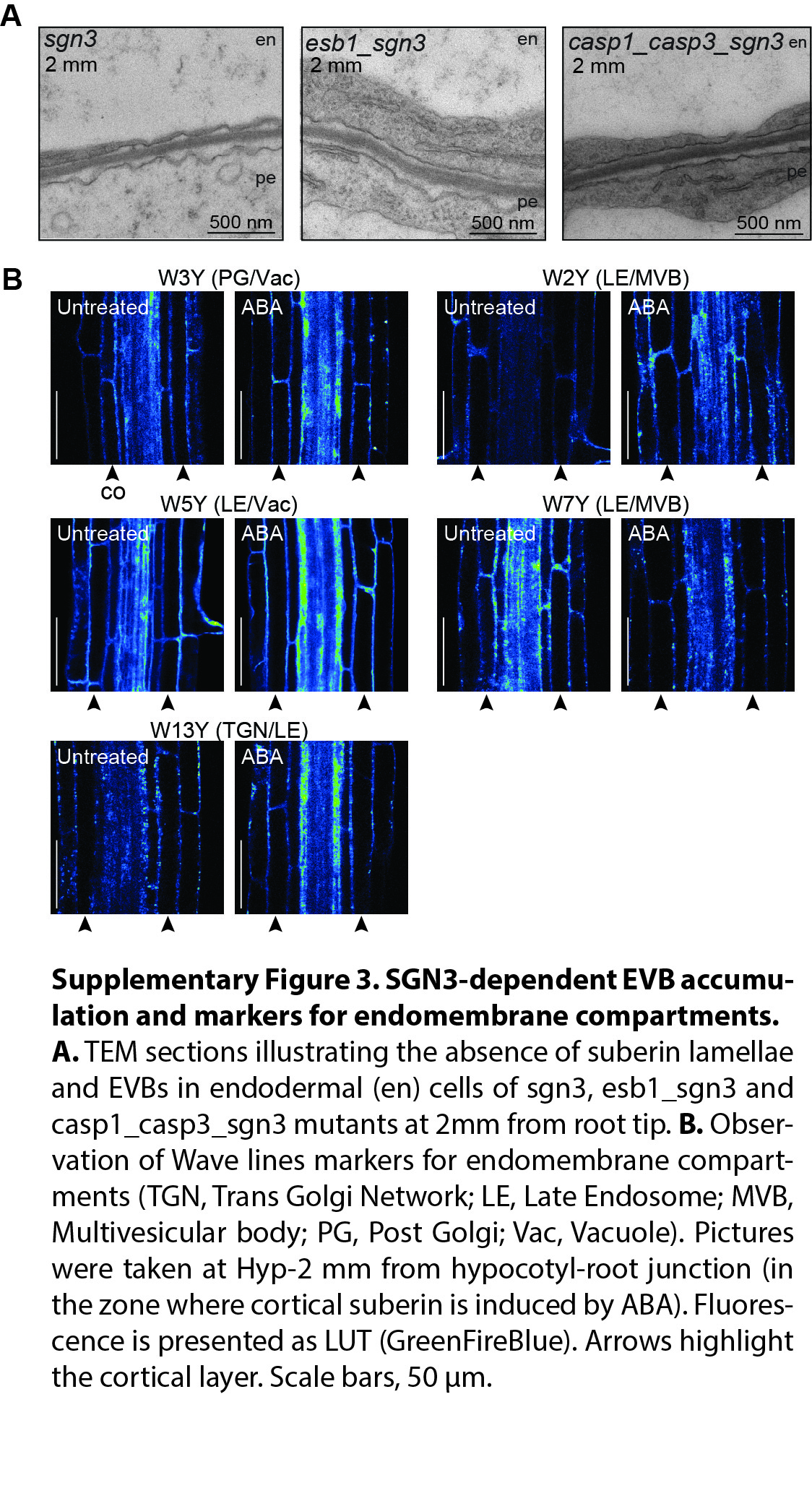
